# Mitigation of Injury from Myocardial Infarction by TH1834, an Inhibitor of the Acetyltransferase Tip60

**DOI:** 10.1101/2021.11.09.467996

**Authors:** Xinrui Wang, Tina C. Wan, Katherine Kulik, Amelia Lauth, Brian C. Smith, John W. Lough, John A. Auchampach

**Affiliations:** Department of Pharmacology and Toxicology, Medical College of Wisconsin, Milwaukee, WI 53226; Department of Cell Biology Neurobiology and Anatomy, Medical College of Wisconsin, Milwaukee, WI 53226; Biochemistry, Medical College of Wisconsin, Milwaukee, WI 53226; Cardiovascular Center, Medical College of Wisconsin, Milwaukee, WI 53226

**Keywords:** Apoptosis, Cardioprotection, Cardiomyocyte Proliferation, Cell-Cycle Activation, Myocardial Infarction, Regeneration, Tip60, TH1834

## Abstract

It is estimated that up to one billion cardiomyocytes (CMs) can be lost during myocardial infarction (MI), which results in contractile dysfunction, adverse ventricular remodeling, and systolic heart failure. Pharmacologic strategies that target factors having both pro-apoptotic and anti-proliferative functions in CMs may be useful for the treatment of ischemic heart disease. One such multifunctional candidate for drug targeting is the acetyltransferase Tip60, which is a member of the MYST family of acetyltransferases known to acetylate both histone and non-histone protein targets that have been shown in cultured cancer cells to promote apoptosis and to initiate the DNA damage response (DDR) thereby limiting cellular expansion. Using a murine model, we recently published findings demonstrating that CM-specific disruption of the *Kat5* gene encoding Tip60 markedly protected against the damaging effects of MI. In the experiments described here, in lieu of genetic targeting, we administered TH1834, an experimental drug designed to specifically inhibit the acetyltransferase domain of Tip60. We report that, similar to the effect of disrupting the *Kat5* gene, daily systemic administration of TH1834 beginning 3 days after induction of MI and continuing for two weeks of a 4-week timeline resulted in improved systolic function assessed by echocardiography, reduced apoptosis and scarring, and increased activation of the CM cell-cycle. Our results support that idea that drugs that inhibit the acetyltransferase activity of Tip60 may be useful agents for the treatment of ischemic heart disease.

## INTRODUCTION

Death of cardiomyocytes (CMs) after prolonged myocardial ischemia results in loss of muscle cell mass, contractile dysfunction, adverse ventricular remodeling, and systolic heart failure. Due to the onset of proliferative senescence that occurs postnatally, CMs are essentially non-regenerable, preventing regeneration of cardiac muscle. Thus, factors that prevent CMs from proliferating and reduce CM survival constitute compelling targets for the therapeutic amelioration of myocardial infarction (MI). We have pursued the lysine acetyltransferase (KAT) termed Tip60 (<u>T</u>at-<u>i</u>nteractive <u>p</u>rotein, <u>60</u> kD) as such a multifunctional candidate. Our rationale is based on findings in the cancer biology field that show that Tip60 acetylates multiple non-histone proteins [1–5] that regulate these functions. For example, Tip60 activates apoptosis by acetylating p53, which in turn trans-activates pro-apoptotic genes [6, 7]. And, Tip60 inhibits cell proliferation by acetylating Atm (Ataxia-telangiectasia mutated [8–10]), thereby initiating the DNA damage response (DDR) culminating in cell-cycle inhibition. We recently reported evidence that Tip60 initiates these activities in CMs at early stages of postnatal heart development [11].

To address the role of Tip60 in the ischemic heart, wherein it is robustly expressed [12, 13], we conditionally disrupted the *Kat5* gene encoding Tip60 in CMs of adult mice three days after induction of MI produced by permanent left coronary artery ligation. As recently reported [14], we found that Cre-mediated depletion of Tip60 markedly preserved cardiac function for up to 28 days post-MI and limited the extent of scarring by ~30%. These findings were accompanied by reduced CM apoptosis, diminished induction of DDR markers in CMs, and activation of the CM cell-cycle. These findings demonstrated that genetic depletion of Tip60 markedly protected against the damaging effects of MI, which might be explained by enhanced CM proliferation and survival [14].

To follow-up these findings in a therapeutic context, we are now addressing whether administration of small molecular weight drugs that inhibit Tip60 acetyl transferase activity produce similar beneficial effects. The experiments reported here describe our initial results examining the novel Tip60 inhibitor TH1834, which was recently developed through in silica modeling efforts of the Tip60 acetyltransferase domain informed by the antimicrobial medication pentamide. Similar to the effect of disrupting the *Kat5* gene, daily intraperitoneal administration of TH1834 in mice from day 3 until day 16 post-MI resulted in improved systolic function throughout the 28-day experimental timeline. Also similar to *Kat5* disruption, TH1834-treated hearts removed at day 28 post-MI exhibited reduced apoptosis and scarring, and increased activation of the CM cell-cycle. These findings warrant further investigation toward the ultimate goal of developing Tip60 inhibitors for the treatment of ischemic heart disease.

## MATERIALS AND METHODS

### Animal Care & Experimentation

These experiments adhered to the National Institutes of Health (NIH) Guide for the Care and Use of Laboratory Animals (NIH Pub. Nos. 85-23, Revised 1996). All protocols described in the authors’ Animal Use Application (AUA #000225), which were approved by the Medical College of Wisconsin’s Institutional Animal Care and Use Committee (IACUC), were adhered to in this study. The IACUC has Animal Welfare Assurance status from the Office of Laboratory Animal Welfare (A3102-01). For these experiments, C57Bl/6 mice were purchased from The Jackson Laboratory.

**Echocardiography** was performed prior to induction of MI or sham surgery and at subsequent intervals using a VisualSonics 3100 high-frequency ultrasound imaging system. This was performed on mice lightly anesthetized with isoflurane delivered via a nose cone (1.0-1.5%). Parasternal long-axis, short-axis, and apical 4-chamber views were obtained using a transducer (MX550D) operating at 30-40 mHz. Short-axis views in M-mode assessed left ventricular anteroposterior internal diameter (LVID), anterior wall thickness (LVAW), and posterior wall thickness (LVPW) at end-diastole (d) and end-systole (s) at the mid-ventricular level. Long-axis views in B-mode assessed left ventricular internal area (LVA) and length (L) at end-diastole and end-systole. Three short axis views (evenly spaced around a mid-level view at the papillary muscles, one towards the apex or distal portion of the heart and one towards the base or proximal portion of the heart) assessed internal area (AreaMid, AreaDist, and AreaProx) at end-diastole and end-systole. Left ventricular systolic function was assessed by the following: (i) fractional shortening (FS %), calculated as ([LVIDd - LVIDs]/LVIDd)*100; (ii) fractional area change (FAC %) = ([LVAd - LVAs]/LVA d)* 100, and (iii) ejection fraction (EF %) = (end-diastolic volume - end-systolic volume)/end-diastolic volume, wherein volumes were estimated by B-mode: 4π/3*L/2*(LVA÷π(L/2) or (AreaMid + AreaDist + AreaProx)*L/3 (Simpson’s method). In addition, global left ventricular function was assessed by calculating the myocardial performance index: MPI = (isovolumic contraction time + isovolumic relaxation time)/ejection time [15–18]. Time intervals were obtained from pulsed Doppler waveforms of mitral valve inflow and aortic valve outflow obtained from apical 4-chamber views.

**Myocardial Infarction** or sham surgery were induced in 10-12-week-old male mice under general anesthesia with isoflurane (1.5-2.0%) and mechanical respiration (model 845, Harvard Apparatus) via an endotracheal tube with room air supplemented with 100% O_2_. Electrocardiograms (ECG; limb lead II configuration) were continuously recorded (Powerlab) via needle electrodes. Rectal temperature was maintained at 37°C using a servo-controlled heating pad. Following anesthesia onset, mice were subcutaneously injected with sustained release meloxicam (4 mg/kg) to manage post-operative pain. A left-sternal thoracotomy was performed to expose the heart, and the pericardium was opened. To target the MI to the lower half of the ventricle, an 8.0 nylon suture was threaded beneath the left main coronary artery at a level below the tip of the left atrium, with the aid of a microscope. Ischemia was induced by tying a permanent suture with a double knot. Coronary occlusion was verified by visually observing blanching of the myocardium distal to the ligature, and by elevation of the ST segment on the ECG. For sham experiments, the sutures were placed as described above but they were not tightened to occlude the artery. The chest wall was closed using polypropylene suture. Recovery was monitored until mice became fully ambulatory.

**Administration of TH1834** was commenced beginning on the third day post-MI/sham surgery via daily intraperitoneal injections of vehicle (1x PBS) to control mice or PBS containing the Tip60 inhibitor TH1834 (10 mg/kg; C33H40N603.2HCl; MW 641.63) to experimental mice; TH1834 was purchased from Axon Medchem (The Netherlands; cat. #2339). Daily injections were performed for 14 consecutive days, i.e. from the third through the sixteenth day post-surgery. On the day before harvest, mice were intraperitoneally injected with BrdU (1 mg). On the day of harvest (10 or 28 days post-MI), mice were euthanized with CO_2_, hearts were removed, and processed for immunostaining (tissue between the apex and ~1 mm above the suture) as described below.

**Terminal deoxynucleotidyl transferase dUTP nick end-labeling (TUNEL)** staining was assessed using the DeadEnd Fluorometric TUNEL System (Promega #G3250) per the manufacturer’s instructions. The total number of TUNEL-positive cells within sections representing the border and remote zones was manually counted while scanning at 400x magnification. TUNEL signal was counted only if confined within a DAPI-positive nucleus, and, nuclei were scored as TUNEL-positive only if at least 50% of the nucleus contained fluorescent signal. Attempts to immunofluorescently co-stain TUNEL-stained sections for markers of CM identity were unsuccessful, presumably due to removal of antigen during proteinase-K digestion. We therefore identified CMs based on cell size (WGA, see below) and autofluorescence signal. Apoptosis was also determined by immunostaining cleaved caspase-3 as described below.

**Myocardial Scarring** was assessed in Masson trichrome-stained transverse sections (4 μm thick) of hearts removed at 0.3 mm intervals along the axis between the apex and ~1 mm above the ligation site. Trichrome-stained sections were photographed at 10x magnification on a Nikon SMZ800 microscope and MIQuant software was used to quantitate infarct size, as previously described [19]. Results are expressed as the average percentage of area, and the midline length, around the left ventricle.

**Wheat Germ Agglutinin (WGA)** staining was performed using Thermo-Fisher #W11263 Alexa Fluor 350 conjugate. Sections mounted on microscope slides were stained with 50 μg/ml WGA in PBS for 10 minutes at room temperature, followed by thorough washing. Images of CMs in transverse orientation were photographed at 400x magnification and processed using ImageJ software to determine numbers of CMs, and numbers of pixels per CM, as respectively indicative of CM density and CM size. Briefly, the DAPI (350) channel displaying CMs outlined in cross-section was isolated, followed by thresholding to fill-in spaces occupied by CM cytoplasm, then adjusting settings to acquire particle sizes in the 600-infinity range having a circularity of 0.25-1. After results (which were set to “include holes”) were obtained, particles representing CMs that were non-transversely sectioned, or blood vessels, were removed.

### Immunostaining & Cell Counting

On the day before harvest, mice were injected with 1 mg 5’-bromo-2’-deoxyuridine (BrdU; Sigma #B9285). Following removal, hearts were perfused with ice-cold cardioplegic solution and atria were removed. Ventricles were fixed overnight in fresh ice-cold 4% paraformaldehyde/PBS, processed through ethanol series, and embedded in paraffin. Sections (4 μm thick) mounted on microscope slides were de-waxed, subjected to antigen retrieval (100°C in 10 mM trisodium citrate pH6.0/0.05% Tween-20 for 20 minutes) followed by 30 minutes’ cooling at RT, and blocked with 2% goat serum/0.1% Triton-X-100 in PBS. Primary antibodies were diluted in blocking buffer and applied overnight at 4°C; secondary antibodies were applied for one hour in the dark. Combinations and dilutions of primary and secondary antibodies used to immunostain each target antigen are listed in Supplemental Table 1.

Microscopy was performed on a Nikon Eclipse 50i microscope equipped with a Nikon DSU3 digital camera. During counting, at least 1,000 CMs were evaluated in 5-6 random 200x photomicrographic fields. CM identity was verified by co-immunostaining cytoplasm with cardiac troponin-T (cTnT), and cellular outline with WGA. CMs in the border and remote (the area ~2 mm distal to the infarct boundary) zones relative to the infarction were separately counted.

### Statistics

All determinations were performed in blinded fashion and are reported as means ± SEM. Echocardiography data were analyzed by a two-way repeated measures ANOVA (time and genotype) to determine whether there was a main effect of time, genotype, or a time-genotype interaction. If global tests showed an effect, post hoc contrasts between baseline and subsequent timepoints within experimental groups were compared by a Dunnett’s multiple comparison *t* test; differences between genotypes at each timepoint were compared by a Student’s *t* test with the Bonferroni correction for multiple comparisons. All other data were compared by an unpaired, two-tailed Student’s t test with Welch’s correction. P < 0.05 was considered statistically significant.

## RESULTS

### TH1834 Experimental Strategy

Our objective was to assess whether the damaging effects of MI could be reduced by subsequent pharmacological inhibition of the acetyltransferase domain of Tip60. As shown in Figure 1A, among commercially developed anti-cancer drugs designed to target KATs, most non-selectively target the similarly structured acetyltransferase domain that is shared by KATs including Tip60 [20]. Curiously, utilization of anacardic acid, MG149, C646, curcumin and garcinol - agents that non-selectively target KATs - to treat cardiac insult has yielded results that are generally positive (Fig. 1A). To our knowledge, pentamidine, NU9056, and TH1834, which are relatively Tip60-specific, have not been utilized for this purpose. We decided to initially interrogate TH1834, based on its structure (Fig. 1B) which was rationally designed to bind a unique pocket within Tip60’s AT domain [20, 21]. As shown in the experimental timeline (Fig. 1C), which is similar to the strategy we previously employed to genetically target *Kat5* [14], we assessed the effect of daily intraperitoneal administration of 10 mg/kg TH1834, beginning on the third day after MI or sham surgery, for 14 consecutive days, ending on the sixteenth day post-MI. During this period, cardiac function was monitored by echocardiography at the indicated intervals (Fig. 1C). On day 10 or 28 post-MI, hearts were removed for histological assessment.

**Figure 1.**
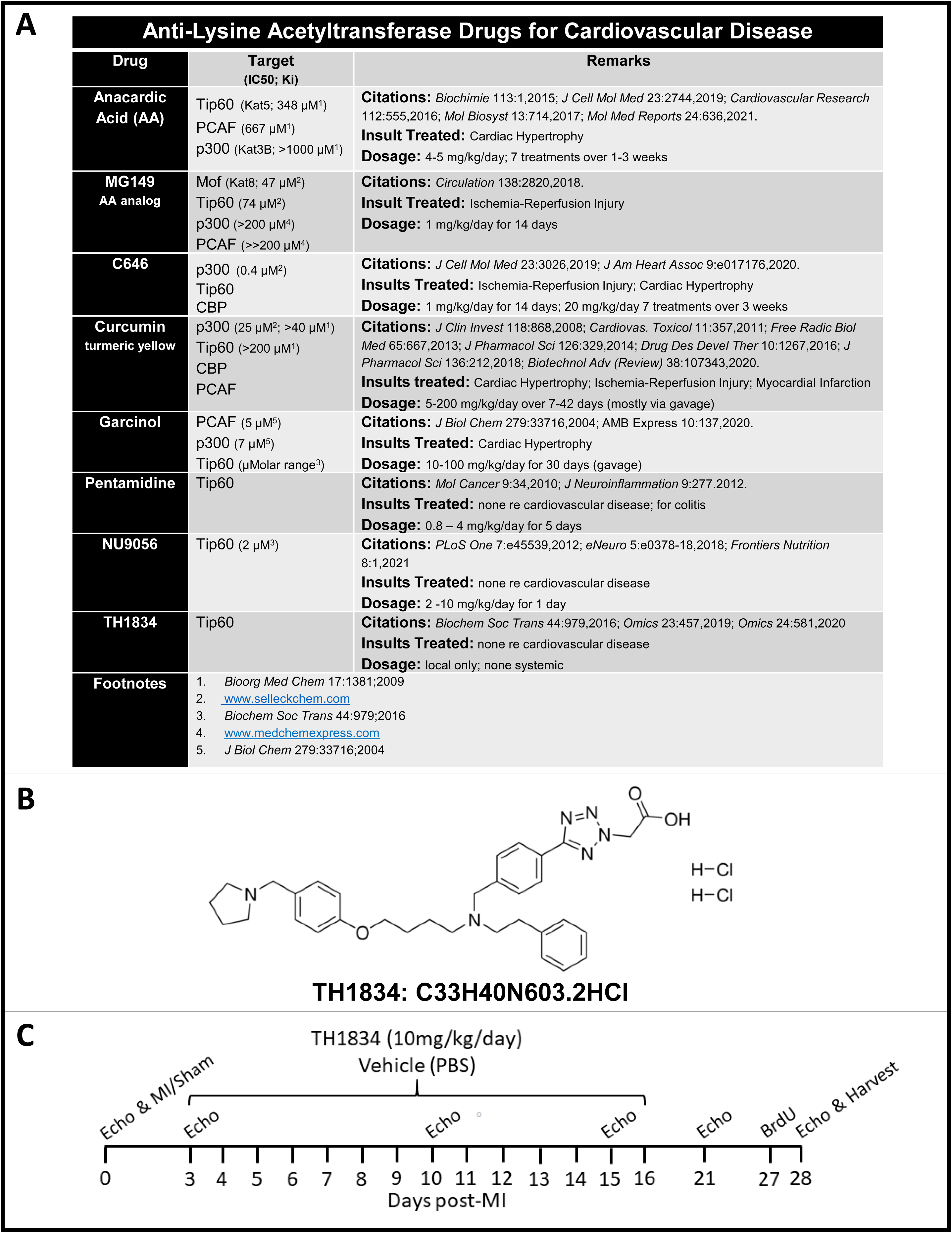
Experimental Strategy: **Panel A**, Tabulated drugs that target KAT acetyltransferase domains. **Panel B**, TH1834 structure. **Panel C**, Experimental Timeline: Hearts of CL57/Bl6 adult mice were infarcted or subjected to sham surgery, followed three days later by the daily intraperitoneal injections of TH1834 (10 mg/kg) for 14 consecutive days.

### Treatment with TH1834 Improves Cardiac Function Post-MI

The mode of cardiac injury employed was designed to generate uniform infarctions in the inferior half of the ventricle by permanently ligating the left main coronary artery below the tip of the left atrium. Per the timeline (Fig. 1C), echocardiography was performed on infarcted mice at intervals up to 28 days post-MI. As shown in Figure 2, in comparison with cardiac function at baseline (BL), dysfunction of all echocardiographic indices (denoted †) was observed at all post-MI timepoints in both control and TH1834-treated mice, with the notable exception that fractional shortening (%FS; Fig. 2B and Suppl Table 2) in TH1834-treated mice was preserved by 10 days post-MI and thereafter. Ejection fraction (EF) calculated by both the B-mode measurements (Suppl Table 3) and Simpson’s method (Fig. 2A and Suppl Table 4) were comparable and were significantly improved by administration of TH1834. Moreover, comparison of control and TH1834-treated mice at each timepoint revealed that, beginning on day 10 post-MI, the TH1834 group exhibited improved Fractional Area Change (FAC), Myocardial Performance Index (MPI), and Stroke Volume (SV), which sustained until the experiment was terminated on day 28 post-MI, when hearts were removed for histology. As shown in Supplemental Figure S1, indices of cardiac function were not altered in TH1834-treated sham-operated mice, indicating that improved function in the MI experiments was not due to a general effect of the drug to enhance cardiac inotropy, but rather must be explained by alternative mechanisms. Details and summaries of echocardiographic data, and heart/body weight ratios, are provided in Supplemental Tables 2-5.

**Figure 2.**
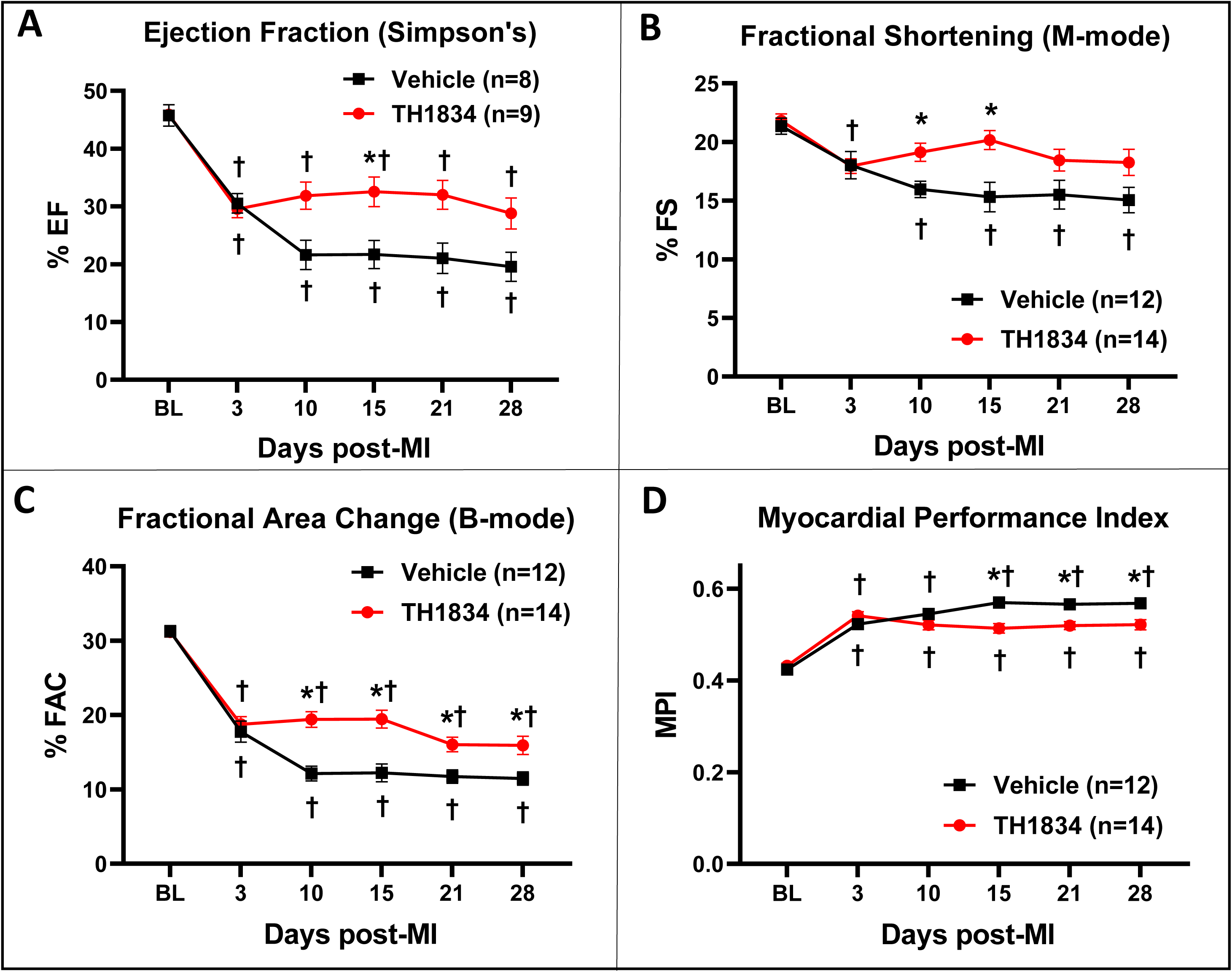
Administration of TH1834 preserves cardiac function after myocardial infarction (MI). Echocardiography was performed at the indicated intervals up to 28 days post-MI, when hearts were removed for histological assessment. **Panels A-D** show indices of LV function, respectively Ejection Fraction (EF), Fractional Shortening (FS), Fractional Area Change (FAC), and Myocardial Performance Index (MPI). Beginning on Day 10, in comparison with vehicle-treated controls, mice treated with TH1834 exhibited improved function. Additional parameters are listed in Supplemental Tables 2-5. Echocardiographic data were analyzed by two-way repeated measures ANOVA followed by Dunnett’s (effect of time) and Bonferroni’s (effect of genotype) multiple comparisons. * denotes P<0.05 vs. Vehicle. † denotes P<0.05 vs baseline (BL) value on Day 0.

### Post-MI Treatment with TH1834 Reduces Scarring

Because Tip60 is pro-apoptotic [6, 7], it was of interest to assess whether treatment with TH1834 on days 3 through 16 post-MI reduced apoptosis, as well as myocardial scarring due to infarction, in hearts removed for histological inactivation at day 10 or 28 post-MI. Apoptosis was assessed by cleaved caspase-3 (Fig. 3A) staining, and by TUNEL (Fig. 3B). Enumeration of caspase-3-positive CMs, as identified by cTnT/WGA staining, revealed significantly reduced numbers at day 10 post-MI. Staining of CM TUNEL-positive nuclei yielded a similar result.

**Figure 3.**
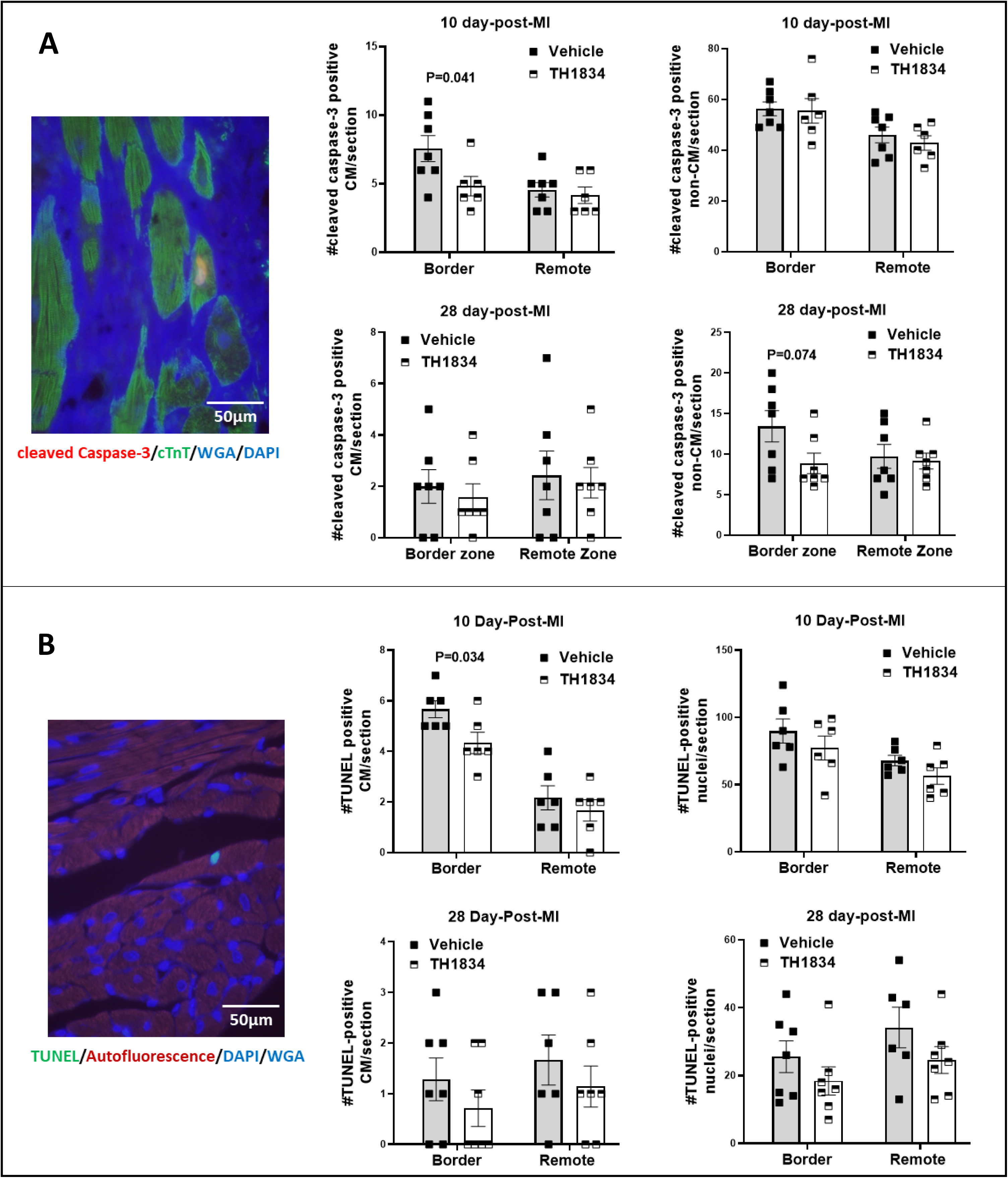
Effects of TH1834 treatment on apoptosis in infarcted mouse hearts. **Panel A**, representative immunostain and quantification of apoptosis per cleaved caspase-3. **Panel B**, representative immunostain and quantification of apoptosis per TUNEL. Data are reported as Mean±SEM and compared by unpaired, two-tailed Student’s t tests with Welch’s correction.

To directly evaluate myocardial damage, extent of scarring was quantitatively assessed by digitizing blue-stained areas, which are indicative of collagen deposition, in trichrome-stained transverse sections removed at 0.3 mm intervals along the axis between that apex and ligation site (Fig. 4, left). In accord with improved function seen by echocardiography, hearts in TH1834-treated mice exhibited significantly diminished scarring at 10 and 28 days post-MI, as indicated by ~25 percent reductions in both the area and midline length parameters (Fig. 4, right).

**Figure 4.**
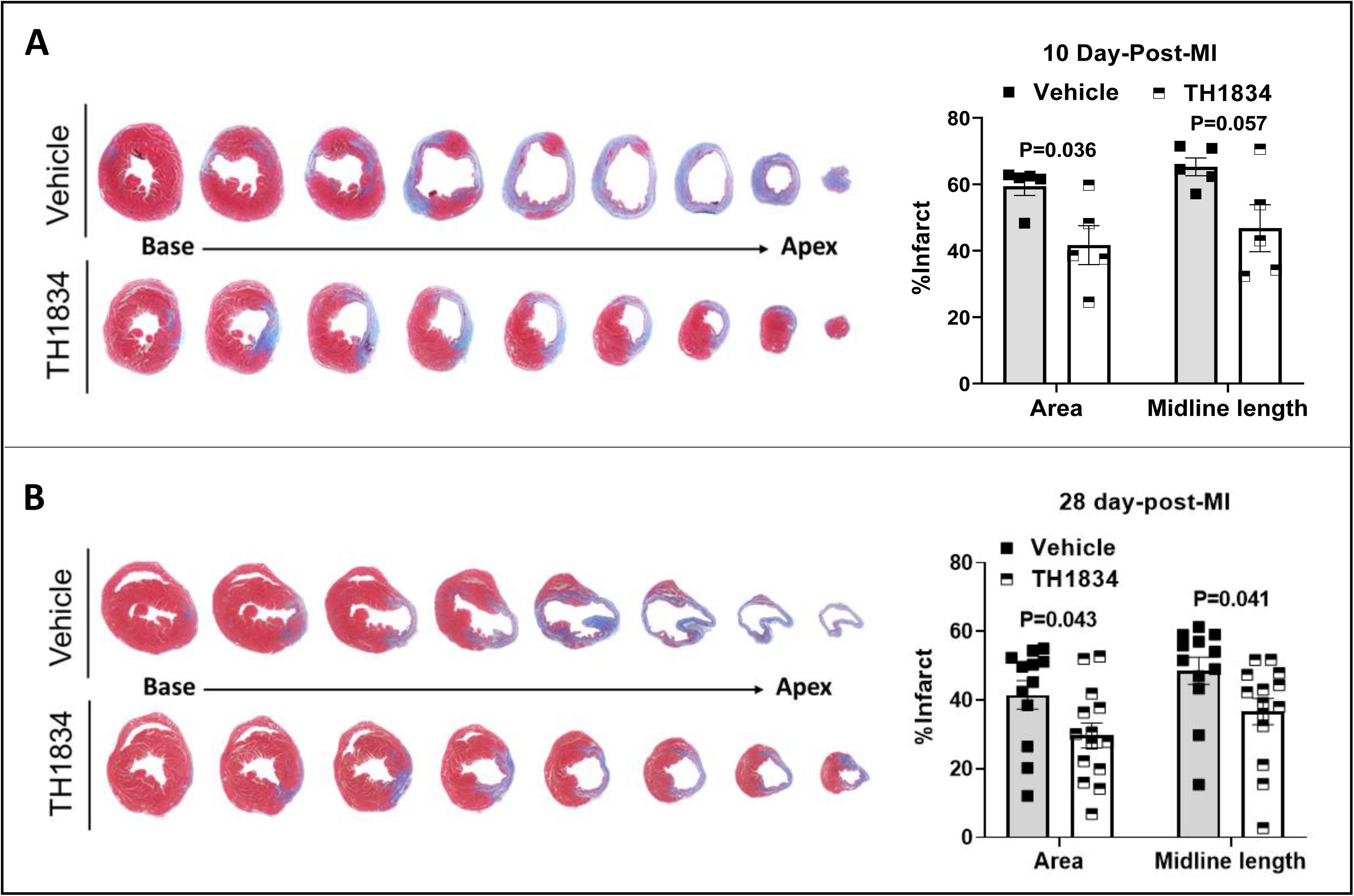
Effects of TH1834 treatment on scarring in infarcted mouse hearts. Representative trichrome-stained cross-sections (left) and scar size quantified by measuring area and midline length below the ligation site (right) at 10 (**Panel A**) and 28 (**Panel B**) days post-MI. Sections were taken at 0.3 mm intervals; blue stain shows area of the scar. Data are reported as Mean ±SEM and compared by unpaired, two-tailed Student’s t tests with Welch’s correction.

### Post-MI Treatment with TH1834 Activates the Cell-Cycle in Cardiomyocytes

To determine whether treatment of mice with TH1834 beginning three days post-MI mimicked the effects of disrupting the *Kat5* gene on cell-cycle activation observed on day 28 post-MI, immunostaining Ki67 (Fig. 5A), BrdU (Fig. 5B), and pHH3 (Fig. 5C) was assessed. These markers respectively identify cells activated throughout the entire cell-cycle, during S-phase only, and in early M-phase. CMs were distinguished from non-CMs by cytoplasmic co-staining with cTnT. As shown in the middle column of each panel of Figure 5, the percentage of CMs exhibiting each of these cell-cycle activation markers was significantly increased in CMs within the border zone of infarcted hearts of mice that had been treated with TH1834, but not with vehicle, at both 10 and 28 days post-MI; trends toward increases in the remote zone were also observed. Similar trends toward cell-cycle activation were observed in non-CMs at day 28 post-MI, ie, hearts in the TH1834-treated group show more Ki67- and BrdU-positive non-CMs, respectively in the border and remote zones (right panels in Fig. 5); although this phenomenon, which is also observed in infarcted hearts following disruption of the *Kat5* gene, is unexplained, we previously speculated that it represents a CM-based paracrine effect, and/or compensated changes resultant from MI [14].

**Figure 5.**
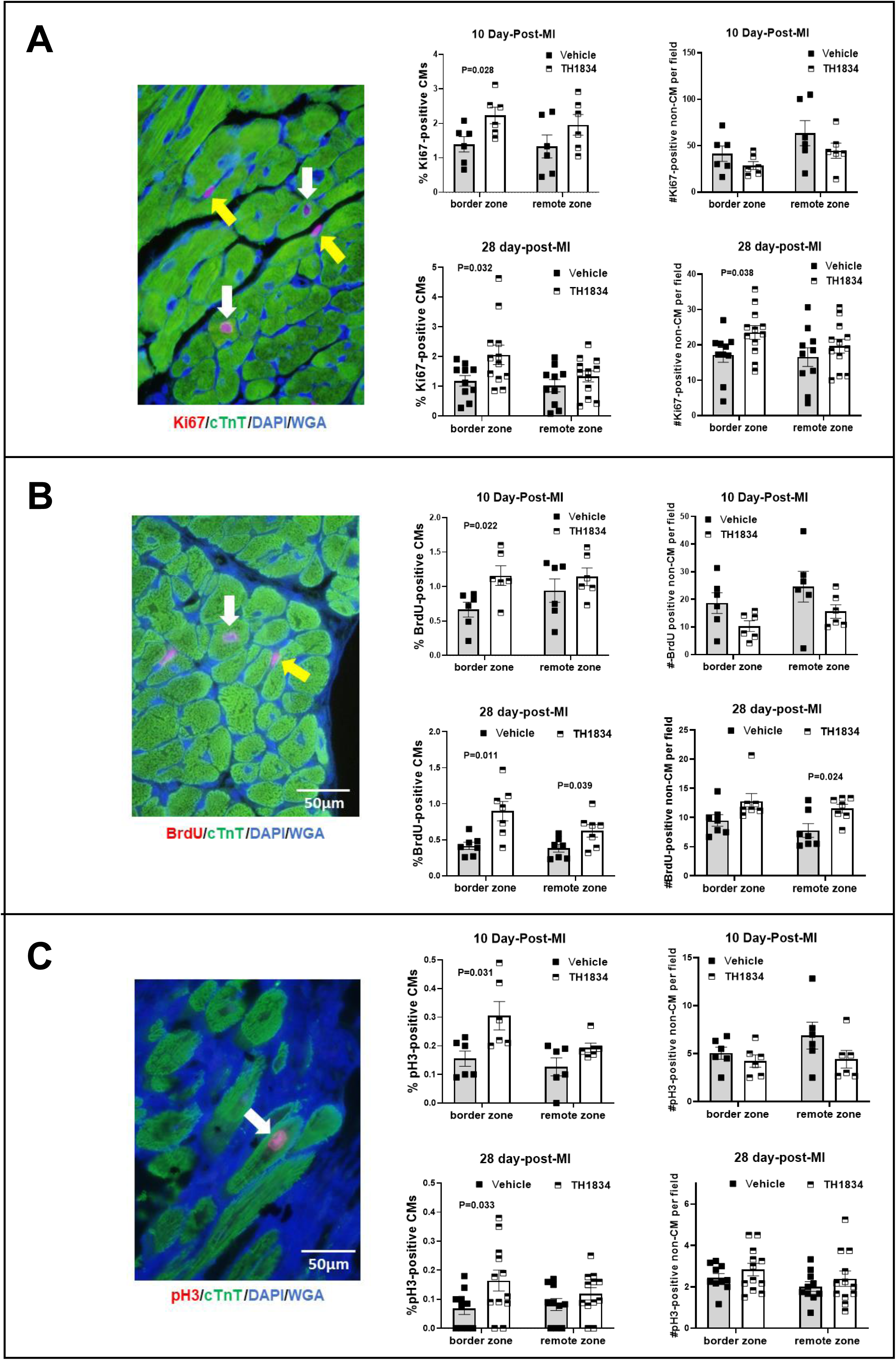
Increased CM cell-cycle activation in infarcted mice treated with TH1834. The **image** of each panel depicts a representative immunostain for each cell-cycle activation marker, denoted by nuclear red fluorescence; each cell was identified as a CM per cytoplasmic expression of cardiac troponin-T (cTnT, green fluorescence). White and yellow arrows respectively denote CMs and non-CMs. The **middle column** of each panel shows percentages of CMs expressing Ki67 (panel A), BrdU (panel B), and pHH3 (panel C) at 10- and 28-days post-MI. The **right column** shows numbers of positive non-CMs per 200x field assessed by evaluating at least 1,000 CMs in six 200x fields per zone in each heart, in blind. Bars denote means ±SEM compared using unpaired two-tailed Student’s t tests with Welch’s correction.

To assess whether CM hypertrophy may have contributed to improved cardiac function in TH1843-treated mice, CM size was estimated by quantitating pixels in transversely sectioned CMs circumscribed by WGA fluorescent staining (Figure 6). This indicated that CM size was not increased in hearts of TH1834-treated mice. In fact, CMs in hearts of mice that had been treated with TH1834 for 14 consecutive days were smaller than in hearts of vehicle-treated mice (Fig. 6, middle column), suggesting that bona fide CM proliferation was induced by inactivating Tip60, a possibility consistent with increased density of CMs in TH1834-treated hearts (Fig. 6 left column).

**Figure 6.**
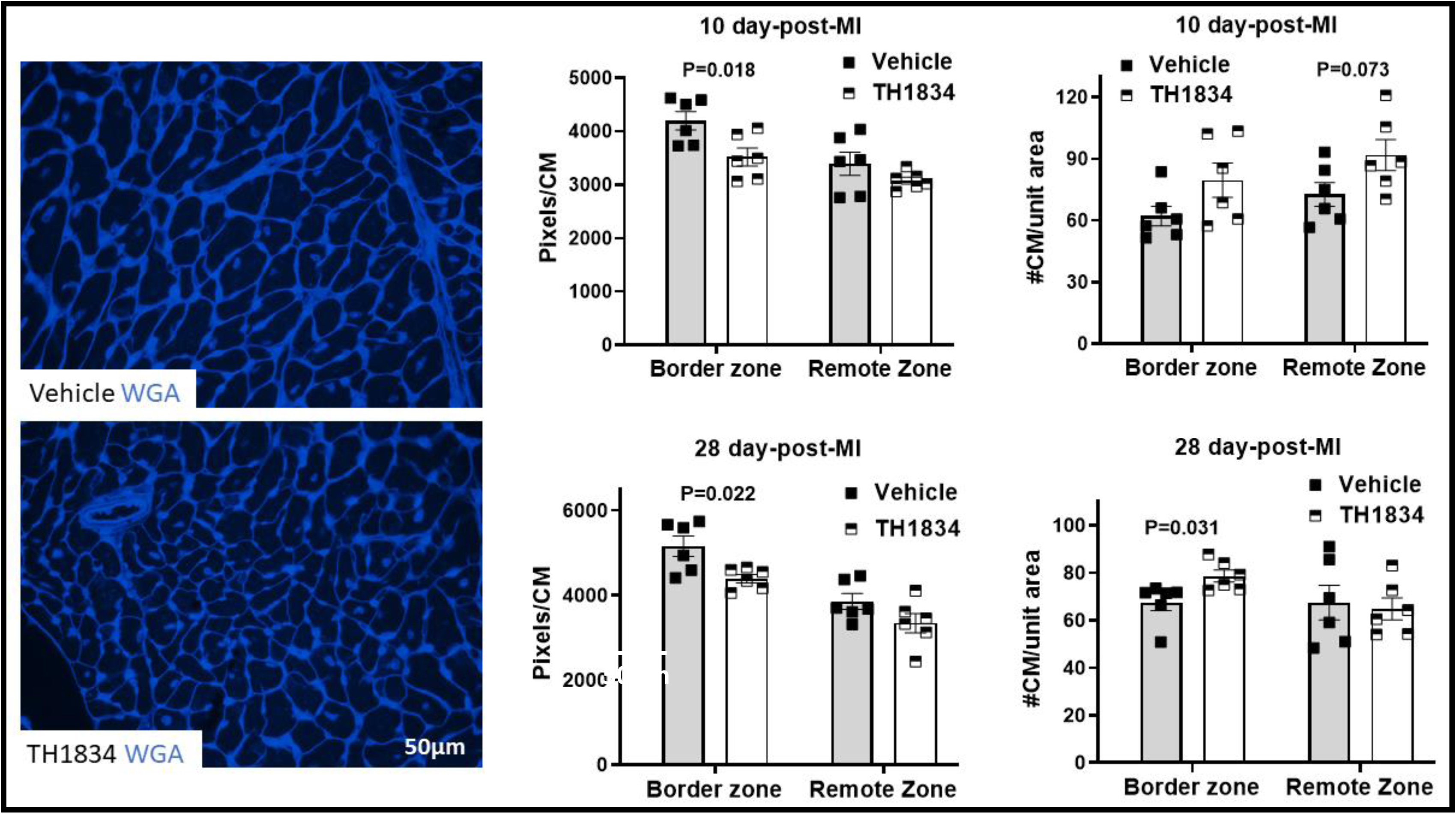
Decreased size and increased density of CMs in the border zone of hearts treated with TH1834. **Left column**, representative areas in the border zone containing transversely-sectioned CMs stained with wheat germ agglutinin (WGA). **Middle column**, bar graphs indicating that CMs in the border zone are smaller (pixels/CM) and **Right column** bar graphs indicating increased in density (#CMs/unit area). Counts were made in blind via ImageJ analysis of >300 transversely-sectioned CMs per zone in each heart. Bars denote means ±SEM compared using unpaired two-tailed Student’s t tests with Welch’s correction.

Finally, because de-differentiation of CMs precedes cytokinesis and proliferation [22], we evaluated gap junction integrity, as indicative of differentiated status, by immunostaining Connexin-43 in hearts of TH1834-treated mice (Supp. Fig. 2). Similar to gap junction dysmorphology we recently described in Tip60-depleted hearts [14], this indicated that the pattern of Connexin-43 staining was disrupted in hearts of the majority of mice treated with TH1834.

## DISCUSSION

The goal of this study was to test the hypothesis that following MI, pharmaceutical inhibition of Tip60, a KAT known to promote apoptosis and inhibit the cell-cycle in cultured cells, protects and regenerates the *in vivo* myocardium by respectively limiting apoptosis and permitting re-activation of the CM cell-cycle. The findings described above are consistent with the possibility that post-MI treatment with a systemically delivered anti-Tip60 drug can diminish scarring while activating the CM cell-cycle, resulting in improved cardiac function.

As recently reviewed, accumulating evidence indicates that the myocardial acetylome is disrupted in various cardiovascular diseases [23]. Although lysine deacetylases (KDACs) and KATs have therefore emerged as therapeutic targets, KATs are receiving less attention [23]. As tabulated in Figure 1A, several drugs are available that inhibit the conserved acetyltransferase domain shared by the ~20 members [24] of the KAT family. Among these, anacardic acid [25–27], MG149 [28], C646 [29, 30], garcinol [31, 32], and curcumin [33–39] - all of which non-selectively target acetyltransferase domains in KATs - have been reported to limit dysfunction caused by cardiac ischemia and hypertrophy. Among these, the polyphenol curcumin, which inhibits p300 (Kat3b; IC_50_ ~40 μM) with reasonable selectivity [40] in comparison with Tip60 (IC_50_ ~200 μM), has received most attention, resulting in improved cardiac function in pre-clinical and clinical studies (review, [39]). And, MG149, which inhibits Tip60 and its closely related family member Mof (Kat8) at respective IC_50s_ of 74 μM and 47 μM, was recently shown to prevent ischemia-reperfusion injury in mice [28]. Considered along with our findings utilizing TH1834, these findings justify expanded efforts to treat cardiovascular disease by targeting individual members of the Kat family with improved specificity.

The mode (intraperitoneal), dosage (10 mg/kg/day), and duration (14 days) selected for deploying TH1834 in this study were estimated based on previous results using the KAT inhibitors cited in Figure 1A and above. In ongoing studies, we are addressing the response of infarcted hearts to various dosages and durations of treatment with TH1834, with emphasis on its effects at earlier timepoints and abbreviated durations of treatment. Also, although we have observed no untoward effects of TH1834, the long-term effects, if any, of transiently inhibiting Tip60, which is considered to possess tumor suppressor function, are being carefully monitored. Studies are also underway to assess whether activation of the CM cell cycle shown in Figure 5 heralds bona fide proliferation and expansion of CM numbers. Finally, alternative to targeting Tip60’s acetyltransferase domain, in order to improve specificity we are designing agents to target its unique chromodomain, which has been shown to be necessary for acetyltransferase function [9].

## List of Non-Standard Abbreviations

Atm: Ataxia-telangiectasia mutated
AurkB: Aurora kinase B
BrdU: 5’-bromodeoxyuridine
CM: cardiomyocyte
cTnT: cardiac troponin-T
Cxn-43: Connexin-43
DDR: DNA damage response
EF: ejection fraction
FAC: fractional area change
FS: fractional shortening
Kat: lysine acetyltransferase
Kat5: lysine acetyltransferase-5 (Tip60)
MI: myocardial infarction
MPI: myocardial performance index
pHH3: phosphohistone H3
Tip60: <u>T</u>at-<u>i</u>nteractive <u>p</u>rotein <u>60</u> kD
TH1834; TUNEL: terminal deoxynucleotidyl transferase dUTP nick end-labeling
WGA: wheat germ agglutinin

## Acknowledgements

Supported by NIH 5R01HL131788 (J.A.A. & JWL), NIH 1S10 OD025038 (J.A.A.), Grant #FP00012308 (J.A.A. & J.W.L.) from the Medical College of Wisconsin Cardiovascular Center, and American Heart Association postdoctoral fellowship #828662 (X.W.).

## Authorship Contributions

J.A.A., J.W.L., and X.W. designed the study. X.W., T.C.W., A.L., and K.K. performed the experimental work and analyzed the data. X.W., J.W.L., and J.A.A. wrote the manuscript and prepared the Figures. All authors reviewed, revised, and approved the final version of the manuscript.

