## Supplemental Data for "Mitigation of Injury from Myocardial Infarction by TH1834, an Inhibitor of the Acetyltransferase Tip60"

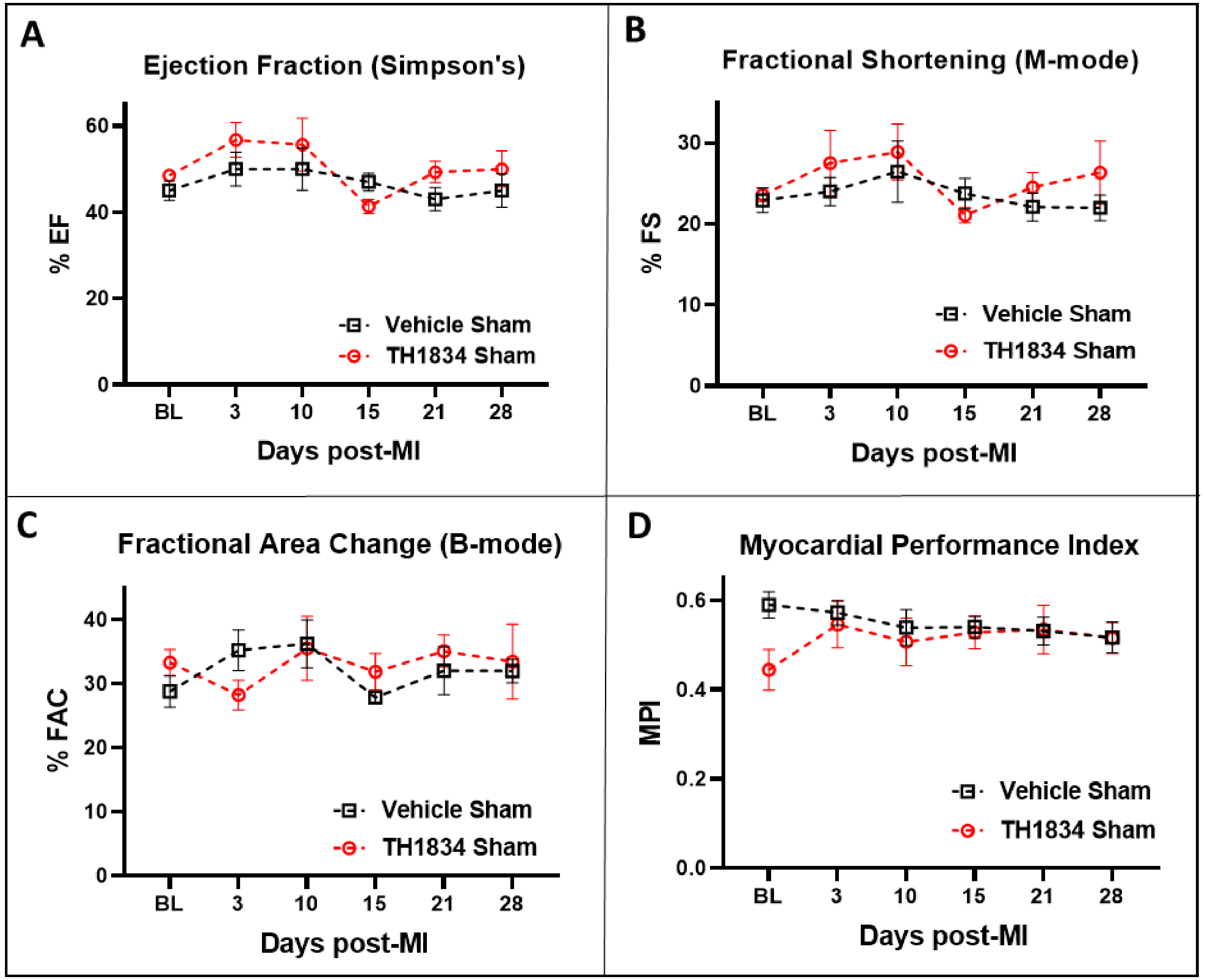

**Supplemental Figure 1. No effect of TH1834 on cardiac function of sham-operated mice.** Echocardiography was performed at the indicated intervals up to 28 days after sham surgery. **Panels A-D** show indices of LV function, respectively Ejection Fraction (EF), Fractional Shortening (FS), Fractional Area Change (FAC), and Myocardial Performance Index (MPI). Echocardiographic data were analyzed by two-way repeated measures ANOVA followed by Dunnett's (effect of time) and Bonferroni's (effect of genotype) multiple comparisons. No significant differences in any of the indices of heart function were identified between the two groups.

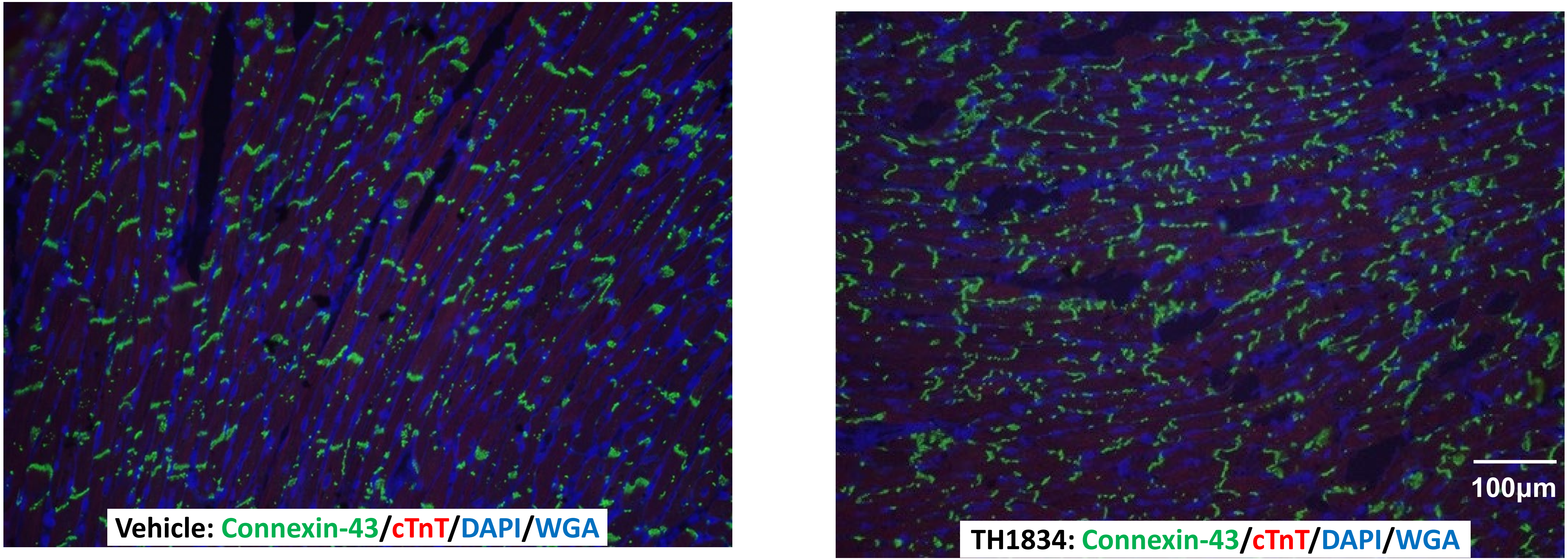

**Supplemental Figure 2**. **Connexin-43 dysmorphology in mice treated with TH1834.** Representative images photographed at 200x magnification showing immunostaining of Connexin-43 in hearts of Vehicle and TH1834 treated mice at 28 days post-MI, indicating that administration of TH1834 after MI is associated with disruption of the intercalated disk. Although not able to be quantified, this phenotype in TH1834-treated hearts was detected in the majority of TH1834-treated hearts by a blinded observer.

### Supplemental Table 1

### Antibodies for Immunofluorescent Staining

| Antigen | Manufacturer | Catalog # | Made in | Dilution |
|---|---|---|---|---|
| 1º 5'-bromodeoxyuridine (BrdU) | Abcam | ab6326 | rat | 1:200 |
| 2º goat anti-rat 594 | Invitrogen | A-11007 | goat | 1:500 |
| 1º caspase-3 | Cell Signaling | 9661S | rabbit | 1:50 |
| 2º goat anti-rabbit 594 | Invitrogen | A-11037 | goat | 1:500 |
| 1º connexin-43 | Abcam | ab11370 | rabbit | 1:1000 |
| 2º goat anti-rabbit 488 | Invitrogen | A-11034 | goat | 1:500 |
| 1º phospho-histone H3 (pH3) | EMD Millipore | 06-570 | rabbit | 1:400 |
| 2º goat anti-rabbit 594 | Invitrogen | A-11037 | goat | 1:500 |
| 1º Ki67 | Invitrogen | 14-5698-82 | rat | 1:250 |
| 2º goat anti-rat 594 | Invitrogen | A-11007 | goat | 1:500 |
| 1º cardiac-Troponin-T (cTnT) | Abcam | ab8295 | mouse | 1:200 |
| 2º goat anti-mouse 488 | Invitrogen | A-11029 | goat | 1:500 |

| | Baseline | | 3 days post-MI | | 10 days post-MI | | 15 days post-MI | | 21 days post-MI | | 28 days post-MI | |
|---|---|---|---|---|---|---|---|---|---|---|---|---|
| **M-Mode & Doppler** | Vehicle N=12 | TH1834 N=14 | Vehicle N=12 | TH1834 N=14 | Vehicle N=12 | TH1834 N=14 | Vehicle N=12 | TH1834 N=14 | Vehicle N=12 | TH1834 N=14 | Vehicle N=12 | TH1834 N=14 |
| LVAW; d (mm) | 0.605±0.029 | 0.645±0.027 | 0.879±0.055† | 0.929±0.048† | 0.830±0.037† | 0.831±0.030† | 0.824±0.024† | 0.825±0.020† | 0.833±0.034† | 0.805±0.027† | 0.768±0.035† | 0.777±0.032† |
| LVAW; s (mm) | 0.791±0.031 | 0.847±0.038 | 1.031±0.058† | 1.086±0.033† | 1.012±0.049† | 1.061±0.033† | 0.977±0.036† | 1.062±0.027† | 0.989±0.032† | 1.025±0.035† | 0.931±0.038† | 0.990±0.040† |
| LVID; d (mm) | 4.319±0.045 | 4.287±0.036 | 3.993±0.098† | 4.056±0.076 | 4.730±0.146 | 4.521±0.113 | 4.910±0.197 | 4.585±0.124 | 5.007±0.210† | 4.818±0.130† | 5.047±0.150† | 4.871±0.176† |
| LVID; s (mm) | 3.397±0.050 | 3.353±0.046 | 3.279±0.110 | 3.334±0.082 | 3.982±0.144† | 3.662±0.111 | 4.182±0.228† | 3.671±0.130 | 4.253±0.238† | 3.943±0.147† | 4.300±0.171† | 4.005±0.193† |
| LVPW; d (mm) | 0.609±0.024 | 0.641±0.013 | 0.806±0.050† | 0.709±0.040 | 0.712±0.062 | 0.779±0.041† | 0.723±0.035 | 0.740±0.042 | 0.745±0.038† | 0.769±0.042† | 0.745±0.040† | 0.754±0.054 |
| LVPW; s (mm) | 0.850±0.021 | 0.884±0.019 | 0.976±0.073 | 0.913±0.043 | 0.867±0.076 | 0.988±0.057 | 0.886±0.045 | 0.914±0.040 | 0.875±0.049 | 0.948±0.045 | 0.946±0.032 | 0.940±0.058 |
| HR (bpm) | 387.3±6.2 | 386.2±9.6 | 442.5±11.2† | 410.1±17.5 | 445.0±16.2 | 441.3±16.4† | 440.0±17.5† | 423.7±15.4 | 433.0±16.0 | 434.5±18.9 | 430.6±19.9 | 444.4±14.9† |
| FS (%) | 21.34±0.68 | 21.81±0.57 | 18.01±1.16 | 17.93±0.62† | 15.95±0.69† | 19.12±0.77* | 15.31±1.25† | 20.16±0.81* | 15.50±1.22† | 18.44±0.92 | 15.04±1.07† | 18.25±1.11 |
| MPI | 0.424±0.003 | 0.432±0.006 | 0.523±0.004† | 0.541±0.009† | 0.545±0.010† | 0.521±0.010† | 0.570±0.012† | 0.514±0.010†* | 0.566±0.011† | 0.520±0.009†* | 0.568±0.011† | 0.521±0.011†* |
| LV mass (mg) | 75.36±4.16 | 79.75±3.11 | 101.7±7.6† | 99.50±4.80† | 119.8±8.9† | 119.0±9.2† | 127.6±7.5† | 117.2±8.9† | 137.5±11.5† | 128.6±10.2† | 131.3±10.2† | 130.4±17.0† |

**Supplemental Table 2: Cardiac Function in Infarcted TH1834-Treated Adult Mice Measured by M-Mode and Doppler Echocardiography.** Ten week-old WT mice were subjected to MI surgery and injected with TH1834 (10mg/kg) or PBS on 14 consecutive days beginning three days after induction of MI. Data are mean ±SEM. LVAW d/s: left ventricle anterior wall thickness in diastole/systole; LVID d/s: left ventricular end diastolic/systolic diameter; LVPW d/s: left ventricle posterior wall thickness in diastole/systole; HR: heart rate; FS: fractional shortening; EF: ejection fraction; MPI: myocardial performance index; LV Vol d/s: left ventricular end-diastolic/systolic volume. *$P<0.05$ vs. Vehicle and †$P<0.05$ vs baseline value analyzed by two-way repeated measures ANOVA followed by Bonferroni's (effect of genotype) and Dunnett's (effect of time) multiple comparisons.

| | Baseline | | 3 days post-MI | | 10 days post-MI | | 15 days post-MI | | 21 days post-MI | | 28 days post-MI | |
|---|---|---|---|---|---|---|---|---|---|---|---|---|
| **B-Mode** | Vehicle N=12 | TH1834 N=14 | Vehicle N=12 | TH1834 N=14 | Vehicle N=12 | TH1834 N=14 | Vehicle N=12 | TH1834 N=14 | Vehicle N=12 | TH1834 N=14 | Vehicle N=12 | TH1834 N=14 |
| Area; d ($mm^2$) | 24.38±0.51 | 24.96±0.53 | 26.07±0.87 | 26.55±0.76 | 33.71±2.40† | 31.91±1.28† | 36.78±2.48† | 31.95±1.13† | 36.16±1.96† | 35.05±2.19† | 37.69±1.72† | 36.37±2.49† |
| Area; s ($mm^2$) | 16.79±0.48 | 17.18±0.39 | 21.47±0.86† | 21.60±0.73† | 29.77±2.35† | 25.81±1.23† | 32.53±2.56† | 25.85±1.21† | 32.05±1.97† | 29.65±2.14† | 33.46±1.76† | 30.92±2.53† |
| FAC (%) | 31.24±0.77 | 31.16±0.65 | 17.76±1.42† | 18.75±1.02† | 12.13±0.99† | 19.41±1.04†* | 12.22±1.19† | 19.45±1.19†* | 11.74±0.89† | 16.03±0.98†* | 11.46±0.88† | 15.91±1.23†* |
| EF (%) | 43.60±1.02 | 44.16±0.78 | 26.38±1.92† | 28.12±1.53† | 18.54±2.40† | 27.37±1.87†* | 16.70±1.87† | 28.53±1.70†* | 17.13±1.74† | 23.21±1.95† | 16.91±1.65† | 23.51±2.09† |
| SV (μL) | 29.43±0.86 | 30.80±0.87 | 19.95±1.43† | 21.96±1.33† | 20.64±1.93† | 28.10±1.33* | 20.65±1.36† | 29.95±1.19* | 21.36±1.46† | 27.06±1.35 | 22.66±1.19† | 28.36±1.12* |
| CO (mL/min) | 11.59±0.30 | 12.06±0.44 | 8.99±0.612† | 9.20±0.56† | 9.62±1.00 | 12.25±0.68 | 10.14±0.82 | 12.68±0.53 | 11.62±1.14 | 12.06±0.95 | 11.17±1.25 | 12.66±0.65 |
| LV Vol; d (μL) | 67.85±2.29 | 69.89±1.95 | 76.76±4.11 | 78.47±3.17 | 124.2±14.3† | 107.7±7.8† | 140.5±15.9† | 109.5±7.0† | 135.4±12.5† | 128.7±14.0† | 143.5±11.6† | 136.9±15.8† |
| LV Vol; s (μL) | 38.42±1.71 | 39.08±1.39 | 56.81±3.76† | 56.51±2.71† | 103.6±14.3† | 79.65±7.39† | 119.9±16.1† | 79.59±6.72† | 114.0±12.5† | 101.6±13.2† | 120.9±11.8† | 108.6±15.3† |

**Supplemental Table 3: Cardiac Function in Infarcted TH1834-Treated Adult Mice Measured by B-Mode Echocardiography.** Ten week-old WT mice were subjected to MI surgery and injected with TH1834 (10mg/kg) or PBS on 14 consecutive days beginning three days after induction of MI. Data are mean ± SEM. Area d/s: left ventricular end-diastolic/systolic area; FAC: fractional area change; FS: fractional shortening; EF: ejection fraction; SV: stroke volume; CO: cardiac output; LV Vol d/s: left ventricular end-diastolic/systolic volume. *$P<0.05$ vs. Vehicle and †$P<0.05$ vs baseline value analyzed by two-way repeated measures ANOVA followed by Bonferroni's (effect of genotype) and Dunnett's (effect of time) multiple comparisons.

| | Baseline | | 3 days post-MI | | 10 days post-MI | | 15 days post-MI | | 21 days post-MI | | 28 days post-MI | |
|---|---|---|---|---|---|---|---|---|---|---|---|---|
| **Simpson's** | Vehicle N=8 | TH1834 N=9 | Vehicle N=8 | TH1834 N=9 | Vehicle N=8 | TH1834 N=9 | Vehicle N=8 | TH1834 N=9 | Vehicle N=8 | TH1834 N=9 | Vehicle N=8 | TH1834 N=9 |
| EF (%) | 45.75±1.85 | 45.90±0.93 | 30.52±1.69† | 29.53±1.49† | 21.59±2.53† | 31.86±2.34† | 21.68±2.43† | 32.54±2.56†* | 21.03±2.63† | 32.01±2.48† | 19.55±2.50† | 28.79±2.68† |
| FAC (%) | 35.70±2.22 | 34.26±1.54 | 24.11±3.05 | 21.40±1.64† | 15.58±3.10† | 28.18±2.79 | 19.18±2.99† | 28.99±2.67 | 15.00±4.17† | 27.46±2.07† | 11.10±3.10† | 24.13±3.52† |
| SV (μL) | 42.44±1.79 | 45.07±1.39 | 32.88±1.95† | 29.83±1.24† | 34.07±2.09 | 44.53±3.05† | 38.46±2.53 | 46.90±3.11† | 36.67±1.52 | 50.24±2.87†* | 35.98±2.12 | 49.84±2.26†* |
| CO (mL/min) | 17.08±0.67 | 17.94±0.61 | 15.05±0.89† | 12.90±0.82† | 14.96±0.99 | 18.66±1.30† | 17.88±1.69 | 19.71±1.39† | 20.36±3.18 | 22.46±1.69† | 16.58±1.54 | 22.28±1.22† |
| LV Vol; d (μL) | 93.25±3.82 | 98.33±2.95 | 108.9±6.8† | 102.1±4.7 | 169.1±17.3† | 143.2±9.5† | 193.3±24.0† | 148.8±10.2† | 192.4±23.5† | 168.2±19.3† | 198.9±19.0† | 187.8±22.1† |
| LV Vol; s (μL) | 50.80±3.10 | 53.25±2.03 | 76.10±5.77† | 72.35±4.47† | 135.0±17.2† | 98.75±9.28† | 154.9±22.8† | 101.9±10.0† | 155.7±23.0† | 117.9±18.4† | 163.0±19.9† | 138.0±21.2† |

**Supplemental Table 4: Echocardiographic Function in Infarcted TH1834-Treated Adult Mice Calculated by Simpson's Method.** Ten week-old WT mice were subjected to MI surgery and injected with TH1834 (10mg/kg) or PBS on 14 consecutive days beginning three days later. Data are mean ±SEM. FAC: fractional area change; FS: fractional shortening; EF: ejection fraction; SV: stroke volume; CO: cardiac output; LV Vol d/s: left ventricular end-diastolic/systolic volume. *P<0.05 vs. Vehicle and †P<0.05 vs baseline value analyzed by two-way repeated measures ANOVA followed by Bonferroni's (effect of genotype) and Dunnett's (effect of time) multiple comparisons.

| | Vehicle<br>N=12 | TH1834<br>N=14 |
|---|---|---|
| Body Weight | 26.63±0.49 | 26.26±0.36 |
| Tibia Length | 16.84±0.05 | 16.88±0.05 |
| Heart Weight | 131.9±3.9 | 126.0±2.7 |
| Heart Weight/Body Weight | 4.95±0.10 | 4.81±0.10 |
| Heart Weight/Tibia Length | 7.83±0.23 | 7.47±0.17 |
| Wet Lung/Dry Lung | 4.03±0.04 | 4.00±0.05 |

**Supplemental Table 5: Heart Weight/Body Weight Ratio 28 Days after Myocardial Infarction (MI).** Ten-week-old WT mice were subjected to MI surgery and injected with TH1834 (10mg/kg) or PBS on 14 consecutive days beginning three days after induction of MI. Data are mean ± SEM.
